# Bioresponsive Microspheres for On-demand Delivery of Anti-inflammatory Cytokines for Osteoarthritis

**DOI:** 10.1101/636886

**Authors:** Eunjae Park, Melanie Hart, Bernd Rolauffs, Jan P. Stegemann, Ramkumar T. Annamalai

**Author notes:** Address correspondence to: Ramkumar T. Annamalai, University of Michigan, 1101 Beal Ave, Ann Arbor, MI 48109.

## Abstract

Despite innovations in surgical interventions, treatment of cartilage injury in osteoarthritic joints remains a challenge due to concomitant inflammation. Obstructing a single dominant inflammatory cytokine have shown remarkable clinical benefits in rheumatoid arthritis, and similar strategies are being suggested to target inflammatory pathways in osteoarthritis (OA). Here we describe the utility of gelatin microspheres that are responsive to arthritic flares, resulting in on-demand, and spatiotemporally controlled release of anti-inflammatory cytokines for cartilage preservation and repair. These microspheres had net negative charge potential to sequester cationic anti-inflammatory cytokines, and the magnitude of the negative charge potential increased with increase in crosslinking density. The enzymatic degradation of the microcarriers was concentration dependent. Release of anti-inflammatory cytokines from the loaded microspheres was directly correlated with the degradation of the gelatin matrix. Exposure of the IL-4 and IL-13 loaded microspheres reduced the inflammation of chondrocytes up to 80%. Hence, the delivery of these microspheres in an osteoarthritic joint can attenuate the stimulation of chondrocytes to secrete catabolic factors including proteinases and nitric oxide. The microsphere format also allows for minimally invasive delivery and is less susceptible to mechanically-induced drug release and are conformant to the intra-articular space. Consequently, bioresponsive microspheres are an effective tool for OA prevention and treatment.

## 1. Introduction

Osteoarthritis (OA) is the primary cause of disability among adults worldwide [1]. OA is a degenerative joint disease that affects one or more diarthrodial joint and is commonly associated with cartilage inflammation and breakdown. OA is a whole-joint disease which includes changes in cartilage, subchondral bone, ligaments, and synovium and subsequent joint failure [2]. OA starts with the “wear and tear” of the cartilage caused primarily due to trauma, obesity, and aging. Although the etiology of OA is still elusive, chondrocyte-mediated inflammatory events driven by the stimulation of innate immune receptors by damage-associated molecules are thought to be the primary cause. The damage-associated molecules activate the synovial macrophages (type A) and fibroblasts (type B) causing synovitis. The activated synovial cells release catabolic and pro-inflammatory factors that lead to a positive-feedback loop of synovial cell activation, inflammation, and cartilage catabolism [3]. When a significant amount of cartilage matrix is degraded, typical symptoms of OA are exhibited, including, joint pain, tenderness, stiffness, loss of flexibility, grating sensation, and bone spurs.

Several therapeutic agents including chondroprotective drugs and growth factors, anti-inflammatory cytokines, extracellular matrix constituents, and immunomodulatory stem cells have been applied to attenuate the cartilage damage [4]. However, the disease-modifying interventions that specifically suppress inflammation are most efficacious for the prevention and treatment of cartilage destruction [5]. Among the many inflammatory cytokines, tumor necrosis factor alpha (TNFα) signaling and interleukin 1 beta (IL-1β) are the predominant cause of cartilage destruction [6]. Although a low level of these factors is necessary for normal homeostasis, inflammatory and oxidative conditions disrupt the balance and drive the pathogenesis of arthritis. Inhibitors of secretion/activity of TNFα and IL-1β including receptor agonists and monoclonal antibodies are shown to mitigate cartilage breakdown [7-9]. More importantly, the application of anti-inflammatory cytokines such as IL-4, IL-10, and IL-13 are shown to reduce inflammation and also to stimulate protective chondrocyte metabolism [4, 10, 11]. Studies have confirmed their inhibitory effects on MMPs secretion, proteoglycans degradation, and chondrocyte apoptosis [12]. Hence, the anti-inflammatory cytokines exhibit both anti-catabolic and anabolic effects for cartilage maintenance. Therefore, these anti-inflammatory cytokines are used as therapeutic agents to modulate the inflammatory response and help prevent chondrocytes hypertrophy and subsequent osteophyte formation [12].

Other therapeutic agents include cartilage anabolic factors such as transforming growth factor beta (TGF-β), fibroblast growth factors (FGF) and insulin-like growth factor-I (IGF-I) that are capable of partially recovering chondrocytes function [4, 13]. Similarly, chondroprotective drugs and matrix constituents such as glucosamine sulfate, chondroitin sulfate, hyaluronic acid, diacerein are shown to decrease NF-kB activation mediated by IL-1β [14]. A key limitation of the anabolic factors and chondroprotective matrices is that they may mitigate the cartilage breakdown but can’t prevent the infiltrated of immune cells. Nevertheless, their effects can be synergistically enhanced when co-delivered with anti-inflammatory cytokines [4]. Alternatively, non-steroidal anti-inflammatory drugs (NSAID) that reduce inflammation by inhibiting the cyclooxygenase enzymes (COX 1 & 2) are commonly prescribed to relieve the joint pain [15, 16]. However, long-term usage of NSAIDs leads to gastrointestinal complications and cardiovascular disease.

Together, delivering anti-inflammatory cytokines to the arthritic joints has the potential to suppress inflammation while stimulating or augmenting chondroprotective metabolism. They are safe, allows quick cartilage recovery, and are effective at almost all stages of the arthritis progression. However, they possess short articular half-lives, which reduce their therapeutic efficacy. For example, IL-10 had a half-life of 2-3 hours in body temperature [17]. Therefore, an efficient drug-delivery vehicle is essential to improve the drug residence time and hence the therapeutic efficacy of anti-inflammatory cytokines.

Here we describe a microsphere drug delivery system that is responsive to arthritic flares, resulting in on-demand and spatiotemporally controlled release of cytokines for cartilage preservation. The microspheres were made out of gelatin crosslinked by genipin, which allows ionic complexation and sequestration of charged anti-inflammatory cytokines. The microspheres are injectable and exhibit degradation and drug release controlled by the proteolytic enzymes characteristically expressed by the inflamed chondrocytes and macrophages in OA. These drug delivery vehicles can titrate the drug release to synchronize with the inflammatory response while reducing the washout of drugs during periods of low disease activity. Based on the endogenous mechanisms, anti-inflammatory cytokines IL-4, IL-10, and IL-13 were chosen for this study. The charger potential of the microspheres to sequester the cationic cytokines was investigated. The degradation of the microspheres and the release kinetics of the cytokines were characterized using enzymatic treatment. Finally, we show that inflammatory cell-mediated release of cytokines from gelatin microspheres has the ability to alleviating the inflammation of activated chondrocytes. Such biomaterial-based approaches can be used to synchronize drug release with the inflammatory response, and thereby prolong the therapeutic effects and the residence time of the cytokines.

**Figure 1.**
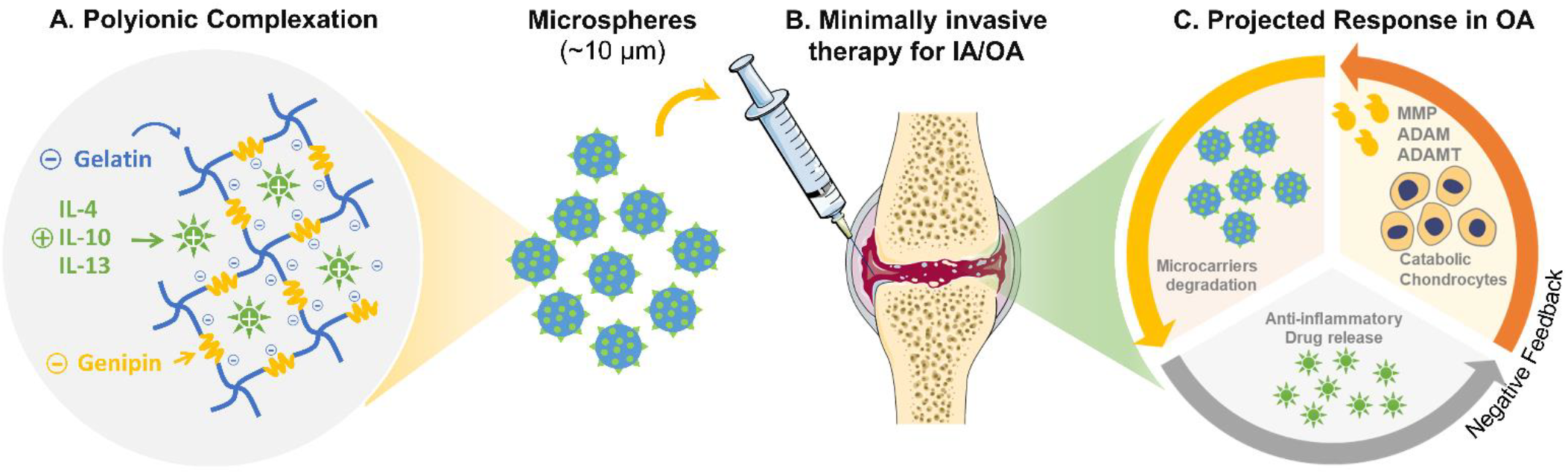
Bioresponsive microspheres for delivery anti-inflammatory cytokines in osteoarthritis. **(A)** The crosslinking of gelatin using amine-reactive genipin imparts a net negative charge on microspheres which is used to sequester cationic cytokines electrostatically. **(B)** The microcarriers allow for minimally invasive delivery and are less susceptible to mechanically-induced drug release and are conformant to the intra-articular space. (C) The microspheres exhibit degradation and drug release controlled by the proteolytic enzymes characteristically expressed by the inflamed chondrocytes and macrophages in OA. These drug delivery vehicles can titrate the drug release to synchronize with the inflammatory response while reducing the washout of drugs during periods of low disease activity.

## 2. Materials and Methods

### 2.1. Microsphere fabrication

Gelatin microspheres were made by emulsification of solubilized gelatin and subsequent crosslinking using genipin. Briefly, gelatin from porcine skin (type A, 175 bloom, Sigma) was dissolved in calcium and magnesium free phosphate buffered saline (1X PBS; Invitrogen, Carlsbad, CA) to make a 6 wt.% stock solution. The stock solution was then dispensed dropwise to polydimethylsiloxane (PDMS) bath kept at 40°C. The mixture was stirred for 5 min using a dual radial-blade impeller at 2000 rpm to emulsify the gelatin solution in PDMS. After the emulsification process, the emulsion was cooled in an ice bath and mixed for an additional 25 minutes to promote gelation. Then 1 wt.% genipin (Wako Chemicals) in PBS was added to the emulsion (2 ml genipin solution per 90 ml emulsion) and mixed at 1500 rpm for additional 30 minutes in the ice bath to initiate cross-linking. The mixture was then collected in 50 ml centrifuge tubes and mixed with 0.1 v/v% of non-ionic surfactants solution and vigorously mixed to encourage phase separation and centrifugation at 200 g for 5 min. After centrifugation, the PDMS supernatant was decanted without disturbing the microspheres pellet. The microspheres were washed three times with the surfactant solution. The influence of surfactants on emulsification was also tested by adding 0.1 v/v% of non-ionic surfactants solution during the emulsification of gelatin in PDMS solution. Two surfactants Pluronic® L101 (denoted as L101, BASF, Milford, CT) and Tween® 20 (denoted as Tween, Sigma) with different hydrophilic-lipophilic balance (HLB) were tested. The microspheres were then suspended in 5 ml of 0.1% genipin in PBS to promote complete cross-linking. The mixture was incubated at room temperature, and samples were collected at 3, 6, 12, and 24 hours and washed with 100% ethanol to remove genipin. The microspheres were then washed three times with deionized water to remove leftover salts and stored in −80°C. Then the frozen microspheres were lyophilized to remove residual water and stored at −20°C for further characterization.

### 2.2. Cell culture

A mouse chondrogenic cell line (ATDC-5, derived from teratocarcinoma AT805, Sigma) and a primary human lung fibroblast cell line (FB, Lonza, Walkersville, MD) obtained from commercial sources were used to characterize gelatin microspheres. For morphological analysis and temporal visualization of cell proliferation using fluorescent microscopy, the FB were transduced with a lentiviral vector expressing green fluorescent protein (GFP) as described previously [18]. Stably transfected cells were selected using puromycin treatment. Finally, highly fluorescent FB were sorted out using cell sorter (MoFlo Astrios EQ, Beckman Coulter) and used for experiments. The chondrogenic cells were cultured in DMEM/F12-Glutamax culture media (ThermoFisher, Waltham, MA) supplemented with 5% fetal bovine serum (FBS, ThermoFisher), and 1% penicillin-streptomycin (ThermoFisher). The fibroblasts were cultured in DMEM media (ThermoFisher) supplemented with 10% FBS and 1% penicillin-streptomycin. The cell response to degradation and cytokine release from the microspheres were characterized by metabolic performance, cell proliferation, and inflammation assays. The cell response was characterized both in 2D (tissue culture treated surface) and 3D pellet cultures (chondrocytes only).

### 2.3. Cell proliferation and viability assay

The cell proliferation was assessed through the quantification of double-stranded DNA using Quant-iT™ Picogreen™ dsDNA assay kit (Invitrogen), and the metabolic performance was quantified using PrestoBlue™ Reagent (Invitrogen). Briefly, cells were seeded at a cell density of 5,000 cells/cm^2^ in a 24-well plate, and after 48 hours of incubation, microspheres were added. To avoid aggregation, 5 mg of microsphere were suspending in 1 ml of PBS and sonicated in ice for 2 minutes at 10% amplitude using a probe sonifier (Branson, Danbury, CT). Then 100 μg of microspheres were added to each well on top of the cells, and cell proliferation and metabolic activity were quantified at day 0, 1, 3 and 7 using corresponding assays. The values were then compared to untreated controls.

### 2.4. Inflammatory cytokine treatment and quantification of inflammation

The capability of drug-loaded microspheres to resurrect chondrocytes under osteoarthritic conditions was tested as follows. Microspheres loaded with IL-4, IL-10, and IL-13 along with corresponding positive and negative control conditions were tested. First, the chondrocytes were seeded at a cell density of 50,000 cells/cm^2^ in a 24-well plate and stimulated with inflammatory cytokines and endotoxin to mimic the inflammatory environment of OA. Several inflammatory cytokines and endotoxin were tested including recombinant murine IL-1β (Peprotech, Rocky Hill, NJ), lipopolysaccharide (LPS, Sigma), interferon gamma (IFNγ, Peprotech), and TNF-α (Peprotech) to induce an inflammatory chondrocyte phenotype. Then the inflammatory state of chondrocytes was quantified by measuring the nitric oxide (NO) production using Griess Reagent Kit (Invitrogen), as described previously [19]. Briefly, 75 µl of culture supernatant was mixed with 75 µl of Griess reagent (1% sulfanilamide in 5% phosphoric acid and 1% N-(1-naphthyl) ethylenediamine dihydrochloride), incubated for 5 minutes at 37°C, and the optical density was measured at 540 nm using a plate reader (Synergy H1™, Biotek, Winooski, VT).

### 2.5. Microspheres zeta potential, loading, degradation, and release

The crosslinking of gelatin using amine-reactive genipin imparts a net negative charge on microspheres and a distinct fluorescence response at 590/620 Ex/Em wavelengths. The zeta potential of the microspheres is the assessment of the charge affinity that can be used to drive the polyionic complexation with the cationic cytokines including IL-4, IL-10, and IL-13. The zeta potential of the microspheres was measured using a Zetasizer (Malvern, Westborough, MA). The microspheres were suspended in a neutral sucrose solution (13 wt.%) as per the manufacturer’s recommendations to avoid aggregation and settling of the microspheres. The microspheres were sonicated for 2 minutes at 10% amplitude in ice using probe sonifier before each measurement.

Lyophilized microspheres were swelled in PBS and loaded with cytokines by incubation in concentrated cytokine stock solution (20-40 µg/ml) overnight at 37ºC. For loading one mg of dry microspheres, 10-20 µL of the cytokine stock solution was supplied and incubated under static conditions. The stock concentrations of IL-4, IL-10, and IL-13 were kept at 40 µg/m, 40 µg/ml and 20 µg/ml respectively, based on the supplier’s recommendation. Loaded microspheres were then washed in PBS containing 10 mg/mL albumin to remove unbound cytokines. The loading efficiency was determined by measuring the concentrations of cytokines in the wash buffer using the respective cytokine ELISA kit (R&D Systems).

For enzymatic degradation and release studies, 100 µg of microspheres were sonicated, loaded with cytokines, and suspended in collagenase solution (Worthington, CLSPA: Activity 1090 Units/mg) and kept at 37ºC until complete degradation. The degradation of the crosslinked microspheres was measured fluorometrically by measuring the fluorescence of the supernatant every 5 minutes for 48 hours at 590/620 Ex/Em. The fluorescence of the crosslinked microspheres also allowed the visualization of their morphology. The cytokines released from the degrading microspheres was measured by ELISA (R&D Systems).

### 2.6. Statistical analysis

All measurements were performed at least in triplicate. Data are presented as mean ± one standard deviation. ANOVA was used for multi-group comparisons, and Student’s t-test with a 95% confidence limit (two-tailed and unequal variance) was used for paired comparisons and the p-values adjusted with Bonferroni correction wherever applicable. Differences with p<0.05 were considered statistically significant.

## 3. Results and Discussion

### 3.1. Microsphere fabrication and influence of surfactant treatment

The bioresponsive microspheres were fabricated by water-in-oil emulsification of gelatin solution which produced spherical droplets on the same size scale as mammalian cells (10-30 µm). Since the sol-gel transition of gelatin is around 30ºC, it is imperative to crosslink the matrix to enhance stability under physiological conditions. Genipin, a less toxic natural crosslinking agent [20], was employed to stabilize the gelatin matrix which yielded insoluble microspheres. The size of the microspheres can be modified through various factors including the speed of emulsification and the viscosity of gelatin and PDMS solutions. The application of surfactants was investigated to attain a homogenous size distribution of microspheres which enables a more predictable degradation and drug release rates. Tween and L101 are nonionic surfactants commonly used in biotechnology and pharmaceutical applications [21, 22]. The amphiphilic surfactant molecules localize to the water-oil interface which is thermodynamically favored [23]. Addition of these surfactants (0.1 %) during gelatin emulsification yielded homogenous microspheres with a narrower size distribution (15-17 µm in diameter) compared to control PBS condition (27-35 µm in diameter, Fig. 2). This effect is due to the stabilization of the droplets by the surfactant molecules that prevent droplet growth (coalescence) [24]. Also, smaller droplets can be formed more easily in the surfactant treated conditions because of lower dynamic interfacial tensions. The size of the microspheres is also crucial in determining the loading capacity of the vehicles since the internal loading volume is proportional to the square of the radius. Although smaller microspheres can limit the drug loading capacity, they can minimize mechanically induced drug release and facilitate smoother articular movements of the knee. The degree of crosslinking of the gelatin matrix, varied by the incubation period in genipin crosslinking solution, had no significant effect on the microsphere size and its distribution (Fig. 2). Overall the surfactant treatment not only reduced the mean size of the microspheres but also narrowed the size distribution of microspheres compared to that of PBS control condition.

**Figure 2.**
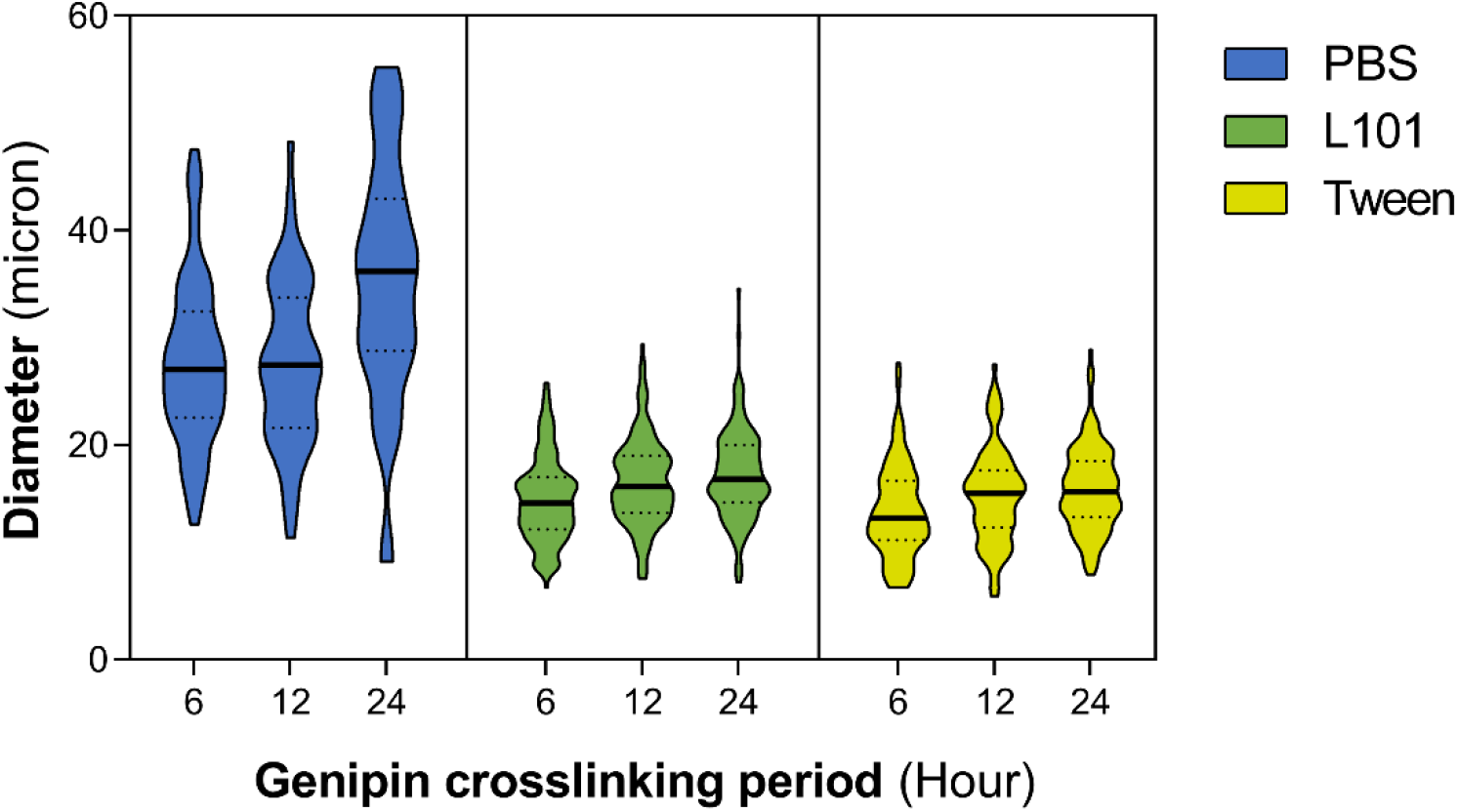
The influence of surfactants on microspheres size distribution. Addition of nonionic surfactants Tween and L101 (0.1 %) during gelatin emulsification yielded homogenous microspheres with a narrower size distribution (15-17 µm in diameter) compared to control PBS condition (27-35 µm in diameter). The degree of crosslinking of the gelatin matrix, varied by the incubation period in genipin crosslinking solution, had no significant effect on the microsphere size and its distribution

### 3.2. Biocompatibility of surfactant treated microspheres

The biocompatibility of the microspheres fabricated through surfactant treatment and genipin cross-linking was investigated through the quantification of cell metabolism and proliferation. Human fibroblasts (FB) were chosen as test cell type and were co-cultured with microspheres fabricated with different treatments. No change in cell metabolism was noticed when the cells were co-cultured with microspheres (Fig. 3A). However, the rate of cell proliferation quantified through DNA assay showed slightly altered growth rates in surfactant-treated conditions. The L101 treatment, although didn’t exhibit any short-term cytotoxicity, it affected the cell-proliferation in the long-term cultures as evident from the ~20% reduction in the cell proliferation at day 7 compared to the control (Fig. 3B). On the other hand, Tween treatment showed short term cytotoxicity as seen from the reduced cell-proliferation rate at 24 hours, but the cells recovered within 7 days of culture (Fig. 3B). The fluorescent micrographs of the GFP-fibroblasts taken at various time points show no differences in cell morphology and proliferation among different conditions (Fig. 3C). The differences in the cytotoxicity of the surfactants are likely due to the differences in hydrophilic-lipophilic balance (HLB) of the surfactants and their subsequent ability to penetrate the cell membrane. Although the effects are minor, Tween was chosen as the standard surfactant for microsphere fabrication due to their long-term biocompatibility. For further characterization, multiple washes were incorporated for thorough removal of residual surfactants from the microspheres.

**Figure 3.**
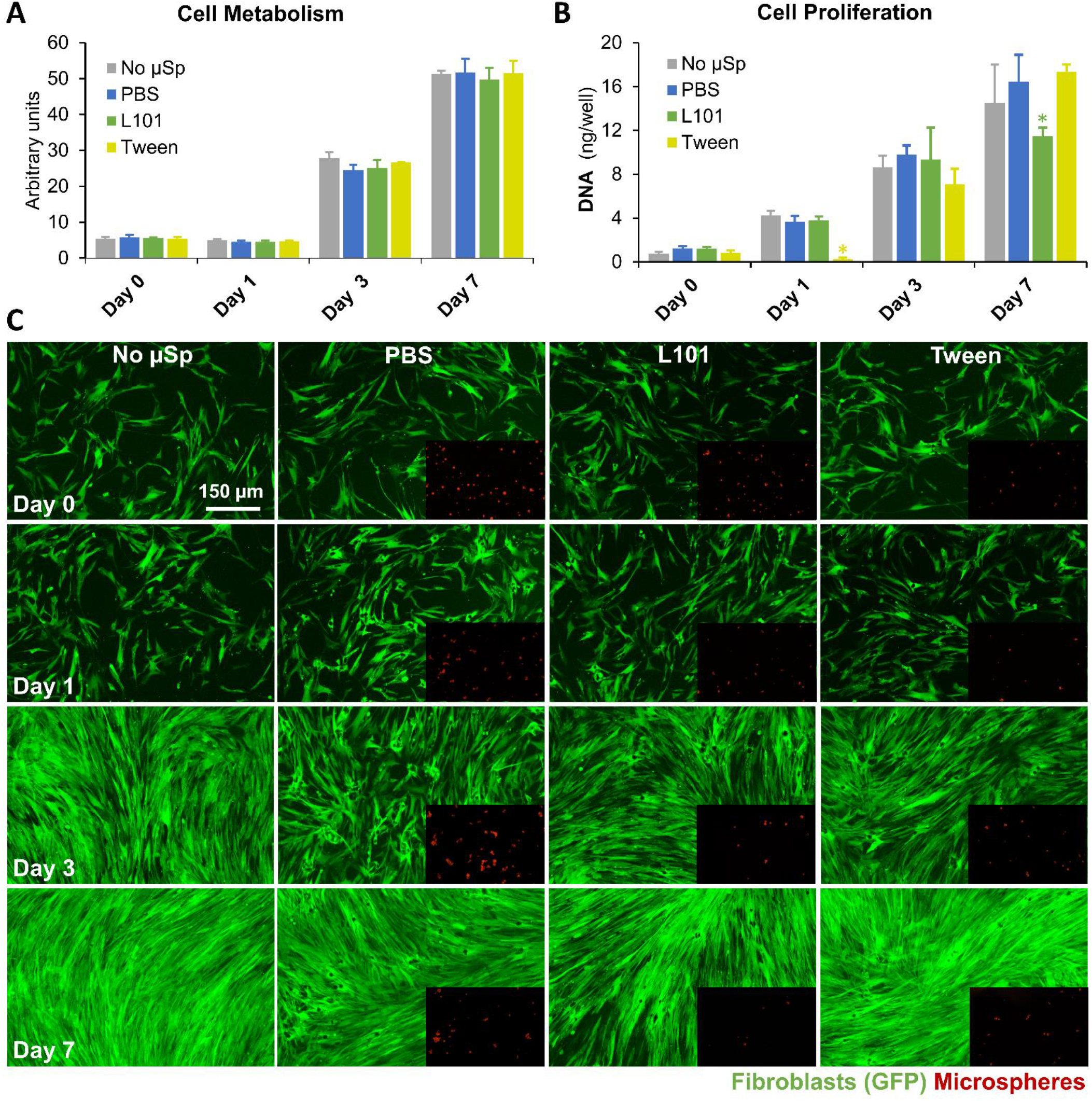
Biocompatibility of the microspheres. (**A**) Human fibroblasts (FB) co-cultured with microspheres fabricated with different treatments exhibited no change in cell metabolism. (**B**) However, the rate of cell proliferation (DNA assay) showed slightly altered growth rates in surfactant-treated conditions. The L101 treatment, although didn’t exhibit any short-term cytotoxicity, a ~20% reduction in the cell proliferation was seen at day 7 compared to the control. Tween treatment, on the other hand, showed short term cytotoxicity, but the cells recovered within 7 days of culture. (**C**) The fluorescent micrographs of the GFP-fibroblasts taken at various time points show no differences in cell morphology and proliferation among different conditions.

### 3.3. Influence of crosslinking density on the charge potential of microspheres

The crosslinking density of the microspheres can be controlled by varying the genipin incubation period [25, 26]. The genipin crosslinking reaction occurs in two steps: First, the primary amines in gelatin form an intermediate through Michael addition, followed by a secondary amide link formation with the genipin ester group through nucleophilic substitution [26]. The influence of the crosslinking density on the charge potential of the microspheres was characterized through zetasizer measurements. Based on the biocompatibility analyses, Tween was chosen as the standard surfactant for microsphere fabrication (Fig. 4A, Confocal images of microspheres fabricated with Tween). The increase in crosslinking density correlated with a decrease in surface charge of the microspheres as measured through the zeta-potential (Fig. 4B). A slight but not significant decrease in zeta-potential was seen between 6-hour and 12-hour incubated samples, but, 24-hour incubated samples showed a drastic decrease of the zeta potential. (p<0.05). The charge potential of the microsphere is an important driving force in ionically complexing the cationic cytokines including IL-4 (pI = 9.17), IL-10 (pI = 8.19), and IL-13 (pI = 8.69) to the gelatin matrix. The control over the crosslinking density and hence the charge potential helps to optimize the cytokines loading and releasing capacity of the microspheres.

**Figure 4.**
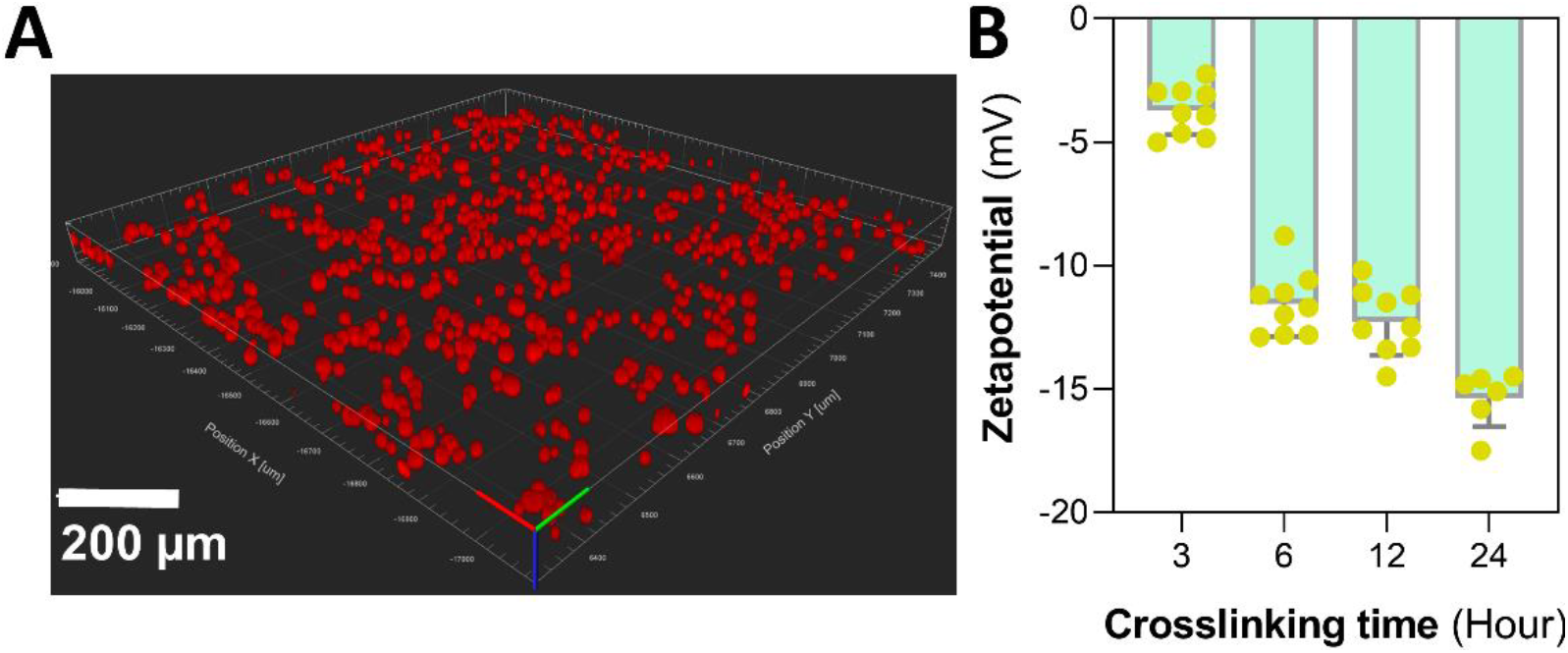
Charge potential of the microspheres. (**A**) Confocal images of microspheres fabricated with Tween treatment showing spherical morphology. (**B**) The zeta-potential measurements of the microspheres showed a correlation between the increase in crosslinking density of gelatin matrix and a decrease in surface charge potential.

### 3.4. Enzymatic degradation and cytokine release from microspheres

The bioresponsivity of the microspheres is the result of their selective degradability to the enzymes characteristically expressed in OA. During inflammation, many contributing cells secrete matrix-degrading catabolic enzymes at levels not usually seen under physiological conditions. These catabolic enzymes can degrade the crosslinked-gelatin matrix in a concentration-dependent manner. The resulting bioresponsivity can be optimized through the method and the extent of crosslinking. In this study, the microspheres were treated with genipin to ensure a crosslinking density of ~90% of the primary amines such that the degradation happens only during the onset of an inflammatory flare.

To demonstrate the bioresponsivity of the microspheres, the microspheres were subjected to enzymatic degradation at increasing collagenase concentrations. Treatment of gelatin microspheres with different concentrations of collagenase at 37°C exhibited a dose-dependent degradation rate (Fig. 5A). The degradation rate was insignificant under control conditions (complete culture media, 0 U/mL of collagenase) and at high collagenase concentrations (>5 U/mL), there was a rapid degradation of the microspheres. At 5 U/mL collagenase treatment, a steady rate of degradation of microspheres was seen that lasted for more than 48 hours. Hence 5 U/mL collagenase treatment was used to characterize the cytokine release from the microspheres. The anti-inflammatory cytokines were loaded into the crosslinked microspheres through electrostatic sequestration by the negatively charged microspheres. The microspheres were loaded with 200-400 ng of IL-4, IL-10, or IL-13 per mg of the microspheres. Upon collagenase degradation, the cytokines were released from the microspheres at rates that linearly correlated with the rate of degradation of the microspheres (Fig.5B).

**Figure 5.**
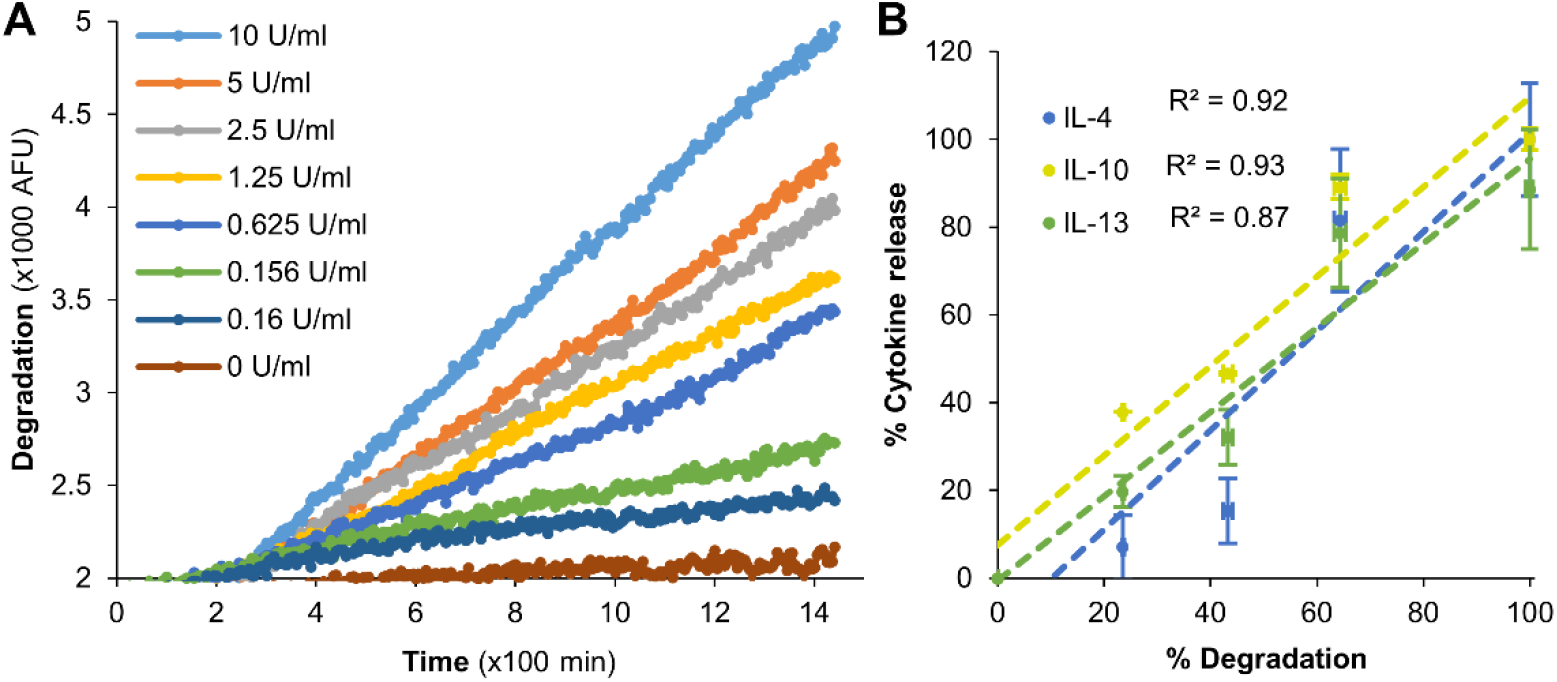
Enzymatic degradation of microspheres and release of cytokines. (**A**) Collagenase treatment of gelatin microspheres at 37°C exhibited a dose-dependent degradation. The degradation rate was insignificant in culture media (0 U/mL of collagenase) while rapid degradation of the microspheres was seen at high collagenase concentrations (>5 U/mL). (**B**) Upon collagenase degradation, the cytokines IL-4, IL-10, and IL-13 were released from the respective microspheres at rates that linearly correlated with the rate of degradation.

We were able to achieve a maximum loading and subsequent release of 40 ng/mg of IL-4 (Fig. 6A), 120 ng/mg of IL-10 (Fig. 6B), 100 ng/mg of IL-13 (Fig. 6C). In all formulations, the release rate of the cytokines linearly correlated with the rate of microsphere degradation (Fig.5, R^2^ ≥ 0.87, p < 0.05). These results demonstrate the on-demand delivery of cytokines from the microspheres in a catabolic microenvironment.

**Figure 6.**
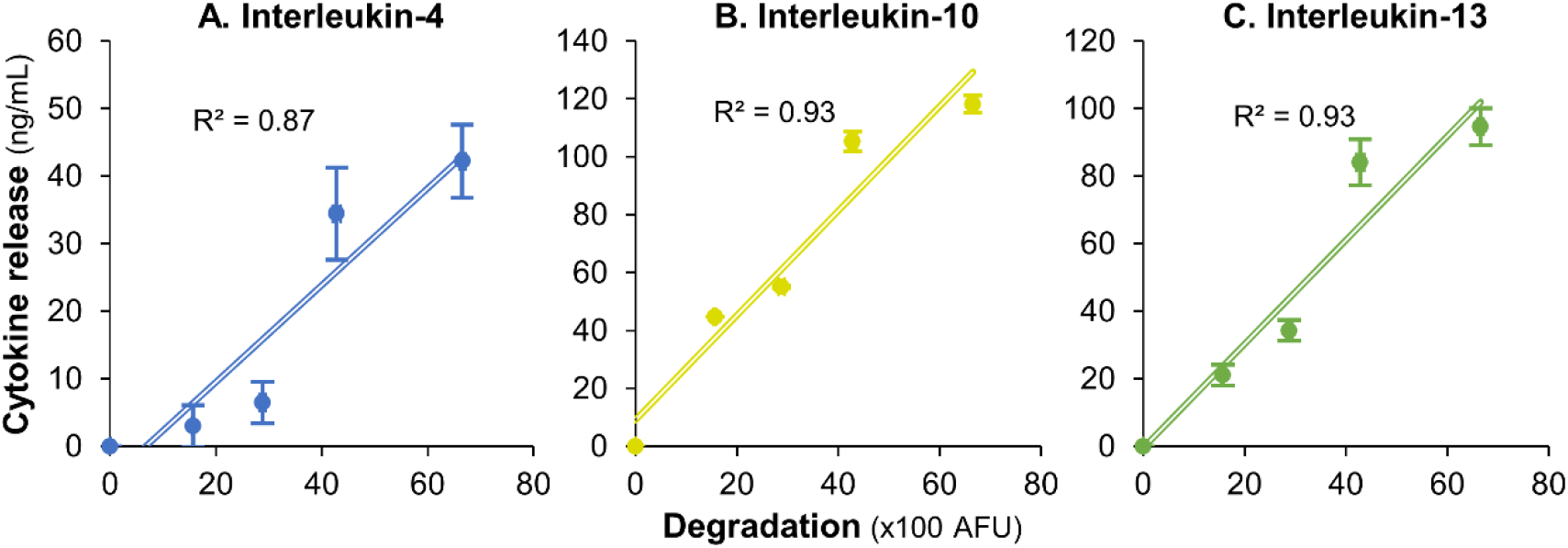
Enzymatic degradation and release of cytokines. A maximum loading and subsequent release of (**A**) 40 ng/mg of IL-4, (**B**) 120 ng/mg of IL-10, and (**C**) 100 ng/mg of IL-13 were achieved using the microspheres. In all conditions, the release rate of the cytokines linearly correlated with the rate of microsphere degradation (R^2^ ≥ 0.87, p < 0.05).

### 3.5. Chondrocyte stimulated with inflammatory agents create an osteoarthritic phenotype

During OA, the anabolic pathways of chondrocytes including the collagen type II and aggrecan production are turned off while the metalloproteinases and nitric oxide production are upregulated [27]. Such an osteoarthritic chondrocyte phenotype would be the ideal culture model to validate the efficacy of drug-loaded microspheres. To create such a phenotype, inflammatory mediators that are commonly implicated in chondrocyte activation were screened. Specifically, lipopolysaccharide (LPS): a toll-like receptor 4 (TLR4) activator, IL-1β, IFNγ, and TNF-α were tested. The resulting osteoarthritic chondrocyte phenotype was evaluated and compared through the quantification of nitric oxide, the key mediator in the progression of OA [28]. Murine IL-1β (2 ng/ml), LPS (200 ng/ml), IFNγ (10 ng/ml), or TNF-α (10 ng/ml) were individually supplemented in the growth media and supplied to the mouse chondrogenic cells (ATDC-5). The cells were cultured for 48 hours, and the cellular nitric oxide (NO) production was measured using Griess assay at various time points (Fig. 7). The assay indicated that TNF-α and IFNγ treatment did not induce any increase in the NO concentration of the cells. However, IL-1β and LPS showed a significant increase in NO production within 12 hours. The chondrocyte activation by IL-1β is mainly induced by the activation of nuclear factor κB (NFκB), which is ubiquitously expressed transcription factor that mediates inflammatory and catabolic event in OA [29]. TLR4, on the other hand, is a type of pattern-recognition receptor that binds not only LPS but also host-derived debris known as damage-associated damage patterns (DAMPs) generated during catabolic events [30]. Therefore, IL-1β and LPS can faithfully replicate the chondrocyte phenotype exhibited in OA. Hence they were chosen for the activation of chondrocyte and subsequent testing of drug-loaded microspheres.

**Figure 7.**
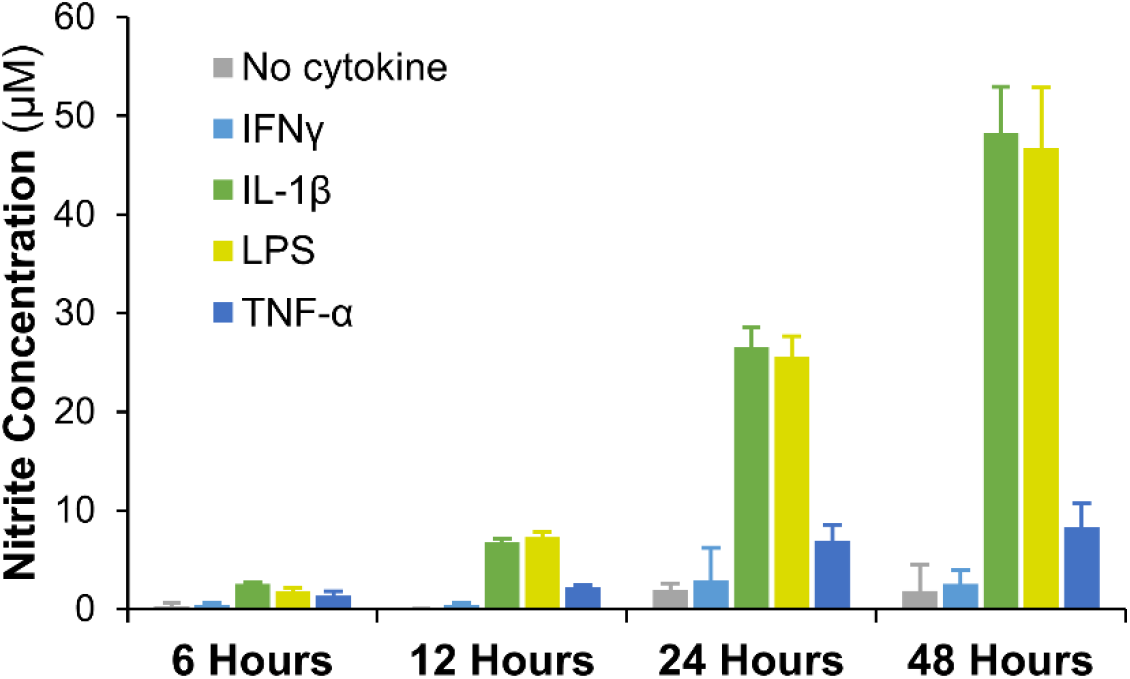
Chondrocyte activation using anti-inflammatory agents. Chondrocytes subjected to TNF-α and IFNγ treatment did not show any increase in the NO production. However, IL-1β and LPS stimulated chondrocytes showed a significant increase in NO production within 12 hours. Therefore, IL-1β and LPS treatment were chosen to replicate the OA chondrocyte phenotype

### 3.6. Microspheres modulated the inflammatory phenotype of activated chondrocytes

Murine chondrocytes (ATDC-5) were stimulated with murine IL-1β (2 ng/ml) or LPS (200 ng/ml) to create an osteoarthritic phenotype that was validated by their upregulated NO production. The activated chondrocytes were then treated with IL-4, IL-10, or IL-13 loaded microspheres (maximum load 40, 120, 100 ng of respective cytokine/mg of microspheres) and the NO production was quantified at day 3. The NO production by the microspheres treated cultures were compared to that of the corresponding bolus treatment cultures in which 200 mg of respective cytokines were directly supplemented in the growth media and replenished every 24 hours (Fig. 8A). The NO production of the chondrocytes showed that the IL-4 and IL-13 released from the microspheres successfully reduced the chondrocyte inflammation by 65-80% within three days (Fig. 8B, Table 1) while IL-10 showed only a marginal effect with no significant decrease in NO production compared to untreated controls. IL-10, although a potent anti-inflammatory cytokine, is known to primarily elicit anti-inflammatory activity only in synergy with other anti-inflammatory cytokines [31]. Hence only a marginal suppression of inflammation is seen in IL-10 treated conditions.

**Figure 8.**
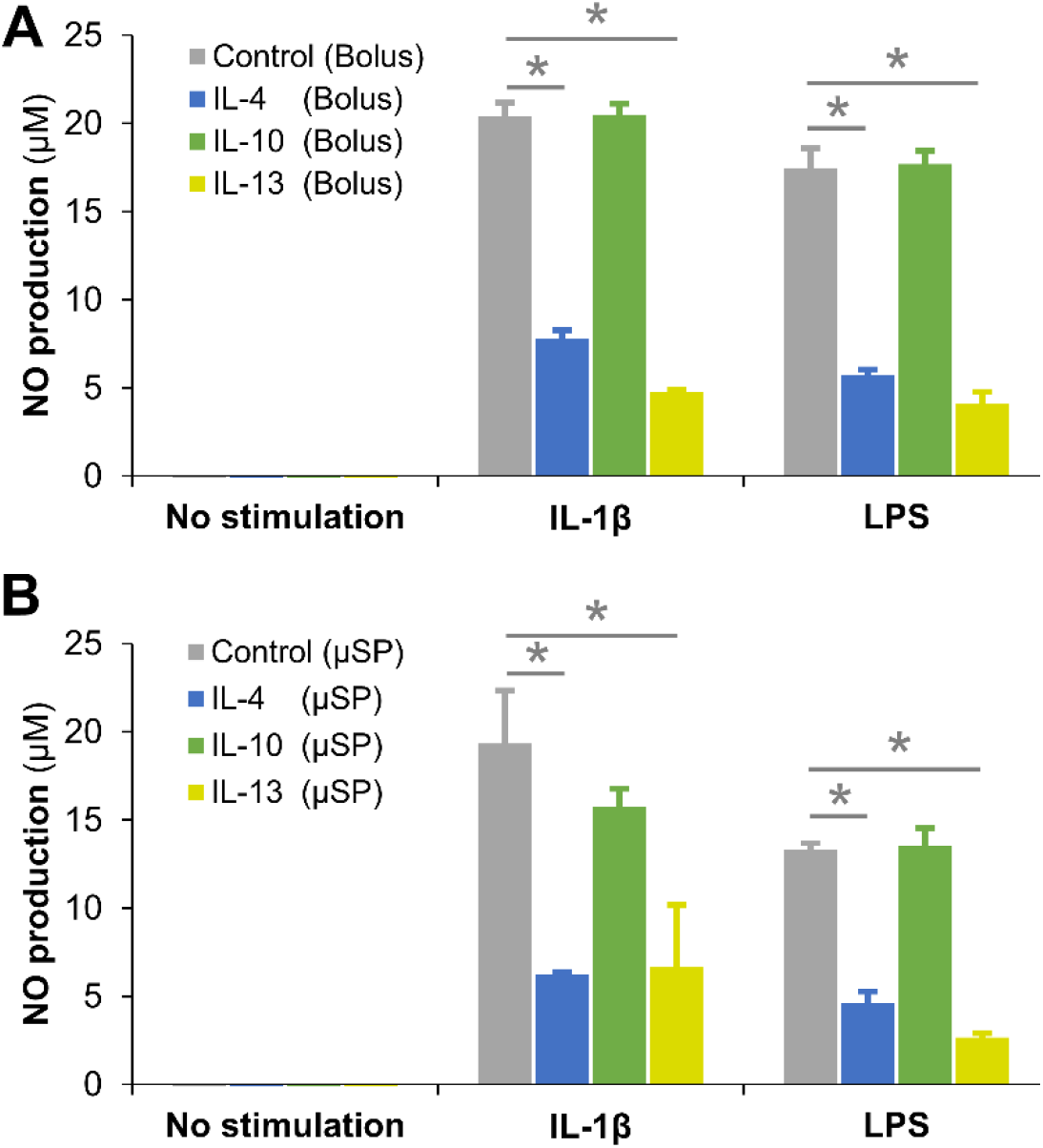
Modulation of inflammation by cytokine loaded microspheres. NO production by the chondrocytes treated with (A) bolus supplementation of 200 mg of cytokines in the culture media and (B) microsphere treatment condition were quantified and compared. The results showed that the IL-4 and IL-13 released from the microspheres successfully reduced the chondrocyte inflammation by 65-80% within three days while IL-10 showed only a marginal effect with no significant decrease in NO production compared to untreated controls. Compared to bolus treatment, the microspheres mediated delivery of the anti-inflammatory cytokines exhibited comparable or better performance.

**Table 1.** Anti-inflammatory cytokine treatment of activated chondrocytes.

|  | Bolus treatment |  | Microsphere treatment |  | Bolus vs. Microsphere |  |
| --- | --- | --- | --- | --- | --- | --- |
|  | *IL-1B* | *LPS* | *IL-1B* | *LPS* | *IL-1B* | *LPS* |
| **IL-4** | 61.83±2.4 | 67.07±1.6 | 67.73±0.7 | 65.36±4.9 | p<0.05 | n.s. |
| **IL-10** | -0.4±3.2 | -1.4±4.3 | 21.01±5.3 | -1.83±7.4 | p<0.05 | n.s. |
| **IL-13** | 76.59±0.7 | 76.39±3.7 | 65.7±18.3 | 80.02±1.8 | n.s. | n.s. |

Compared to bolus treatment, the microspheres mediated delivery of the anti-inflammatory cytokines exhibited comparable or better performance (Table 1). It should be noted that 2-5 times more cytokines were added to the bolus treatments compared to the microsphere treatment and were replenished every 24 hours to account for the short half-lives of the cytokines. These results show that the microspheres are efficient in titrating the drug release depending on the inflammatory response while reducing drug washout. Further, the microspheres are biodegradable and can be quickly removed from the synovium once digested by the catabolic factors. Their breakdown products are non-toxic and didn’t show any agonistic influence on inflammatory pathways. This is noteworthy since the extended residence of non-digestible substances is known to cause inflammation and contribute to OA. Further, microcarriers format allows for minimally invasive delivery and is less susceptible to mechanically-induced drug release and are conformant to the intra-articular space. Taken together, bioresponsive microspheres are an effective tool for osteoarthritic prevention and treatment.

## 4. Conclusion

We have created bioresponsive gelatin microspheres that can prolong the half-life of anti-inflammatory cytokines while reducing their washout during periods of low disease activity. The gelatin microspheres are made through a facile emulsification procedure followed by appropriate post-processing that allowed tailoring the crosslinking density and the charge potential of the matrix. The negative charge potential of the microspheres allowed ionic complexation and sequestration of cationic anti-inflammatory cytokines. We previously showed that the genipin crosslinking density could be optimized such that the microspheres can be preferentially degraded by inflammatory cells and not by other non-inflammatory cells [26]. Applying the same principle, the microspheres were made preferentially degradable by the catabolic factors symptomatically secreted by the inflamed chondrocytes and macrophages in OA. The enzymatic degradation of the microspheres was concentration dependent and the release of the cytokines linearly correlated with the degradation rates. The IL-4 and IL-13 loaded microspheres co-cultured with osteoarthritic chondrocytes reduced their inflammation by up to 80%. Such on-demand delivery systems that are synchronized with the catabolic responses may find general utility in wound healing particularly in preventing inflammation-mediated cartilage damage in OA.

## Reference

1. Neogi, T., The epidemiology and impact of pain in osteoarthritis. Osteoarthritis and cartilage, 2013. 21(9): p. 1145–1153.

2. Martel-Pelletier, J., et al., Osteoarthritis. Nature Reviews Disease Primers, 2016. 2: p. 16072.

3. Nefla, M., et al., The danger from within: alarmins in arthritis. Nat Rev Rheumatol, 2016. 12(11): p. 669–683.

4. Ying, Z., P. Tyler, and P. Ming, Anti-Inflammatory Strategies in Cartilage Repair. Tissue Engineering Part B: Reviews, 2014. 20(6): p. 655–668.

5. Sokolove, J. and C.M. Lepus, Role of inflammation in the pathogenesis of osteoarthritis: latest findings and interpretations. Therapeutic advances in musculoskeletal disease, 2013. 5(2): p. 77–94.

6. Malemud, C.J., Cytokines as Therapeutic Targets for Osteoarthritis. BioDrugs, 2004. 18(1): p. 23–35.

7. Martel-Pelletier, J., et al., In vitro effects of diacerhein and rhein on interleukin 1 and tumor necrosis factor-alpha systems in human osteoarthritic synovium and chondrocytes. J Rheumatol, 1998. 25(4): p. 753–62.

8. Scott, D.L. and G.H. Kingsley, Tumor necrosis factor inhibitors for rheumatoid arthritis. N Engl J Med, 2006. 355(7): p. 704–12.

9. Kawaguchi, A., et al., Blocking of tumor necrosis factor activity promotes natural repair of osteochondral defects in rabbit knee. Acta Orthop, 2009. 80(5): p. 606–11.

10. Lubberts, E., et al., Regulatory role of interleukin 10 in joint inflammation and cartilage destruction in murine streptococcal cell wall (SCW) arthritis. More therapeutic benefit with IL-4/IL-10 combination therapy than with IL-10 treatment alone. Cytokine, 1998. 10(5): p. 361–9.

11. Lubberts, E., et al., Adenoviral vector-mediated overexpression of IL-4 in the knee joint of mice with collagen-induced arthritis prevents cartilage destruction. J Immunol, 1999. 163(8): p. 4546–56.

12. Wojdasiewicz, P., Ł.A. Poniatowski, and D. Szukiewicz, The role of inflammatory and anti-inflammatory cytokines in the pathogenesis of osteoarthritis. Mediators of inflammation, 2014. 2014: p. 561459–561459.

13. Andia, I. and N. Maffulli, Platelet-rich plasma for managing pain and inflammation in osteoarthritis. Nature Reviews Rheumatology, 2013. 9: p. 721.

14. Gouze, J.N., et al., Glucosamine modulates IL-1-induced activation of rat chondrocytes at a receptor level, and by inhibiting the NF-κB pathway. FEBS Letters, 2002. 510(3): p. 166–170.

15. Noble, S.L., D.S. King, and J.I. Olutade, Cyclooxygenase-2 enzyme inhibitors: place in therapy. Am Fam Physician, 2000. 61(12): p. 3669–76.

16. Jiang, D., et al., Efficacy of intra-articular injection of celecoxib in a rabbit model of osteoarthritis. Int J Mol Sci, 2010. 11(10): p. 4106–13.

17. Suga, M., S. Keshavjee, and M. Liu, Instability of cytokines at body temperature. The Journal of Heart and Lung Transplantation, 2005. 24(4): p. 504–505.

18. Annamalai, R.T., et al., Vascular Network Formation by Human Microvascular Endothelial Cells in Modular Fibrin Microtissues. ACS Biomaterials Science & Engineering, 2016. 2(11): p. 1914–1925.

19. Hayashi, T., et al., Nitric oxide production by superficial and deep articular chondrocytes. Arthritis Rheum, 1997. 40(2): p. 261–9.

20. Tsai, C.C., et al., In vitro evaluation of the genotoxicity of a naturally occurring crosslinking agent (genipin) for biologic tissue fixation. J Biomed Mater Res, 2000. 52(1): p. 58–65.

21. Tomlinson, A., et al., Polysorbate 20 Degradation in Biopharmaceutical Formulations: Quantification of Free Fatty Acids, Characterization of Particulates, and Insights into the Degradation Mechanism. Molecular Pharmaceutics, 2015. 12(11): p. 3805–3815.

22. Gates, K.A., et al., A new bioerodible polymer insert for the controlled release of metronidazole. Pharm Res, 1994. 11(11): p. 1605–9.

23. Lawrence, M.J. and G.D. Rees, Microemulsion-based media as novel drug delivery systems. Advanced Drug Delivery Reviews, 2000. 45(1): p. 89–121.

24. Rowe, E.L., Effect of Emulsifier Concentration and Type on the Particle Size Distribution of Emulsions. J Pharm Sci, 1965. 54: p. 260–4.

25. Solorio, L., et al., Gelatin microspheres crosslinked with genipin for local delivery of growth factors. J Tissue Eng Regen Med, 2010. 4(7): p. 514–23.

26. Annamalai, R.T., et al., Harnessing macrophage-mediated degradation of gelatin microspheres for spatiotemporal control of BMP2 release. Biomaterials, 2018. 161: p. 216–227.

27. Rahmati, M., A. Mobasheri, and M. Mozafari, Inflammatory mediators in osteoarthritis: A critical review of the state-of-the-art, current prospects, and future challenges. Bone, 2016. 85: p. 81–90.

28. Studer, R., et al., Nitric oxide in osteoarthritis. Osteoarthritis and Cartilage, 1999. 7(4): p. 377–379.

29. Kenneth, B.M., et al., NF-κB Signaling: Multiple Angles to Target OA. Current Drug Targets, 2010. 11(5): p. 599–613.

30. Gómez, R., et al., TLR4 signalling in osteoarthritis—finding targets for candidate DMOADs. Nature Reviews Rheumatology, 2014. 11: p. 159.

31. van Roon, J.A.G., F.P.J.G. Lafeber, and J.W.J. Bijlsma, Synergistic activity of interleukin-4 and interleukin-10 in suppression of inflammation and joint destruction in rheumatoid arthritis. Arthritis & Rheumatism, 2001. 44(1): p. 3–12.

